# Dissection of demethylation and toxicity induced gene expression changes after decitabine treatment

**DOI:** 10.1101/2019.12.18.880633

**Authors:** A Turchinovich, HM Surowy, AG Tonevitsky, B Burwinkel

## Abstract

The DNA methyltransferase inhibitor decitabine (DAC) is a widely used drug for both fundamental epigenetics studies and anti-cancer therapy. Besides DNA demethylation, DAC also induces cell toxicity associated with DNA damage. The dual-mode of DAC action on cells provides a significant hurdle to study genes which expression is regulated by CpG methylation. In this work, we performed the analysis of global DNA methylation levels in cultured cancer cells after treatment with increasing doses of DAC and have found the U-shaped curve of the de-methylation efficacy induced by the drug. Specifically, high doses of DAC induced significantly lower DNA hypomethylation as compared to hundred-fold less concentrated doses. At the same time, the impact of DAC on cell viability was dose-dependent. These findings allowed dissecting the demethylation and the cell toxicity impact of DAC on gene expression in subsequent mRNA-seq experiments. Surprisingly, the number of genes that were upregulated due to DNA hypomethylation was comparable to the number of genes induced by DAC toxicity. Furthermore, we show that high DAC concentrations induce downregulation of housekeeping genes which are most widely used for RT-qPCR normalization (including GAPDH, actin and tubulin). The latter suggests that genes unaffected by DAC treatment would manifest themselves as upregulated when their expression is normalized on a downregulated housekeeping reference. Finally, we show that expression of most human oncogenes and tumor-suppressor genes remains unaffected after DAC treatment, and only a few of them were upregulated due to DNA hypomethylation. Our work stresses the importance of closely studying the correlation of the degree of DNA demethylation induced by varying doses of DAC with changes in gene expression levels to avoid erroneous conclusions regarding epigenetic silencing of a gene.

## Introduction

The inhibitor of DNMT methyltransferase 5-Aza-2′-deoxycytidine (decitabine, DAC) have been widely used in multiple epigenetic research studies, and also approved for the treatment of some types of cancers [Kantarjian et al, 2006]. The DAC resides incorporated into the genomic DNA during replication are recognized by mammalian DNA methyltransferase 1 (DNMT1) enzyme that remains covalently bound to the nucleoside, thereby depleting the cells of enzyme activity and resulting in DNA demethylation [Christman 2002; Ghoshal et al, 2005; Schermelleh et al, 2005]. Despite its utilization in numerous fundamental and applied medical studies, the precise mode of DAC action remains poorly understood [Stresemann et al, 2008]. The depletion of DNMT that results in global DNA demethylation is one of the well-confirmed effects of DAC. However, incorporation of DAC into replicating DNA results in formation of covalent DNA-DNMT adducts [Schermelleh et al, 2005], DNA damage and subsequent induction of genes involved in DNA repair, growth arrest and apoptosis [Karpf et al, 2001; Stresemann and Lyko, 2008; Jiemjit et al, 2008; Palii et al, 2008; Ruiz-Magana et al, 2012]. This dual-mode of DAC action necessitates experimental protocols that allow dissecting the hypomethylation- and the toxicity-induced gene expression changes.

In the present study, we used Next-Generation bisulfite sequencing (BS-Seq) to evaluate global CpG methylation in cultured U2OS cells upon treatment with increasing concentrations of DAC. Our results confirmed the previously reported U-shaped curve of DAC-induced hypomethylation efficacy. Specifically, ultra-low (0.01 μM) and high (10 μM) DAC doses mediated significantly lower CpG hypomethylation as compared to low (0.1 μM) drug doses. At the same time, the impact of DAC on cell viability was concentration-dependent. Based on these findings we have defined a set of criteria to differentiate hypomethylation- and toxicity-induced gene expression changes after DAC treatment observed in the subsequent high-throughput mRNA-seq experiments.

Our results are in accordance with previous studies demonstrating that while DAC causes global hypomethylation throughout the entire genome, only a small percentage of genes that undergo promoter hypomethylation becomes reactivated [Tsai et al. 2012; Klco et al. 2013; Pandiyan et al. 2013; Lund et al. 2014]. Dissection of the demethylation and the toxicity related responses of gene expression after DAC treatment could help to support therapeutic decision making and selecting optimal drug concentration for a given type of cancer. Finally, we noticed that housekeeping reference genes that are most widely used for qPCR normalization were significantly downregulated following DAC treatment indicating that caution should be taken when applying qPCR-based methods for the analysis of DAC-induced changes in gene expression.

## Results

### The impact of the increasing doses of DAC on global DNA methylation level and cell viability

The original aim of this study was to investigate the impact of DAC treatment on DNA methylation, viability and gene expression in cultured U2OS cells (Fig. 1A). We initially examined the effects of three different DAC concentrations (ultra-low 0.01 μM, low 0.1 μM, and high 10 μM) on global DNA methylation using whole-genome bisulfite sequencing performed with previously described Capture and Amplification by Tailing and Switching (CATS) protocol [Turchinovich et al, 2014] (Fig. 1B). For that purpose U2OS cells in culture were treated with the indicated DAC doses or DMSO (control cells) for 96 hours (Fig. 1A). Afterward, total DNA was isolated, bisulfite-converted and sequenced on Illumina HiSeq platform after generating CATS bisulfite-DNA libraries (as described in the Material and Methods section). The QC reports for CATS bisulfite sequencing data are available in the Supplementary material (Supplementary Fig.1). Treatment with a low 0.1 μM DAC dose resulted in an approximately 67.5% decrease of CpG methylation as compared to DMSO treated cells (53.50% vs. 17.40% methylated cytosines in CpG context) across the whole genome. As was expected, ultra-low 0.01 μM DAC dose induced a significantly smaller DNA demethylation as compared to 0.1 μM dose (41.30% vs. 17.40% methylated cytosines in CpG context). Surprisingly, DNA from cells treated with high (10 μM) DAC concentrations contained 33.00% methylated cytosines in CpG context. Therefore, 10 μM DAC induced almost two-fold weaker CpG hypomethylation as compared to 100-fold more diluted 0.1 μM DAC dose, and only about 25% higher hypomethylation as compared to ultra-low 0.1 μM DAC (Fig.1B). These results accord with a previous report by Qin and co-authors who have demonstrated the U-shaped hypomethylation dose-response curve of DAC treatment in various cultured cells [Qin et al, 2007]. The optimal concentrations of DAC were cell type-dependent but lied within the 0.05 μM – 0.5 μM range for most cell types [Qin et al, 2007]. Lower demethylating efficacy of high DAC concentrations was also explained by a more significant interference of the drug in cell proliferation, which is essential for the incorporation of DAC into the DNA. Indeed, the U2OS cells treated with 10 μM DAC exhibited a more impaired cell growth rate as compared to low and ultra-low doses (Fig.3C). Thus, after 96h of incubation the number of viable cells in 10 μM DAC treated wells was approximately two-fold lower as compared to control wells, while the number of cells in 0.1 μM DAC wells was only about 25% lower than in controls (Fig.3C).

**Figure 1.**
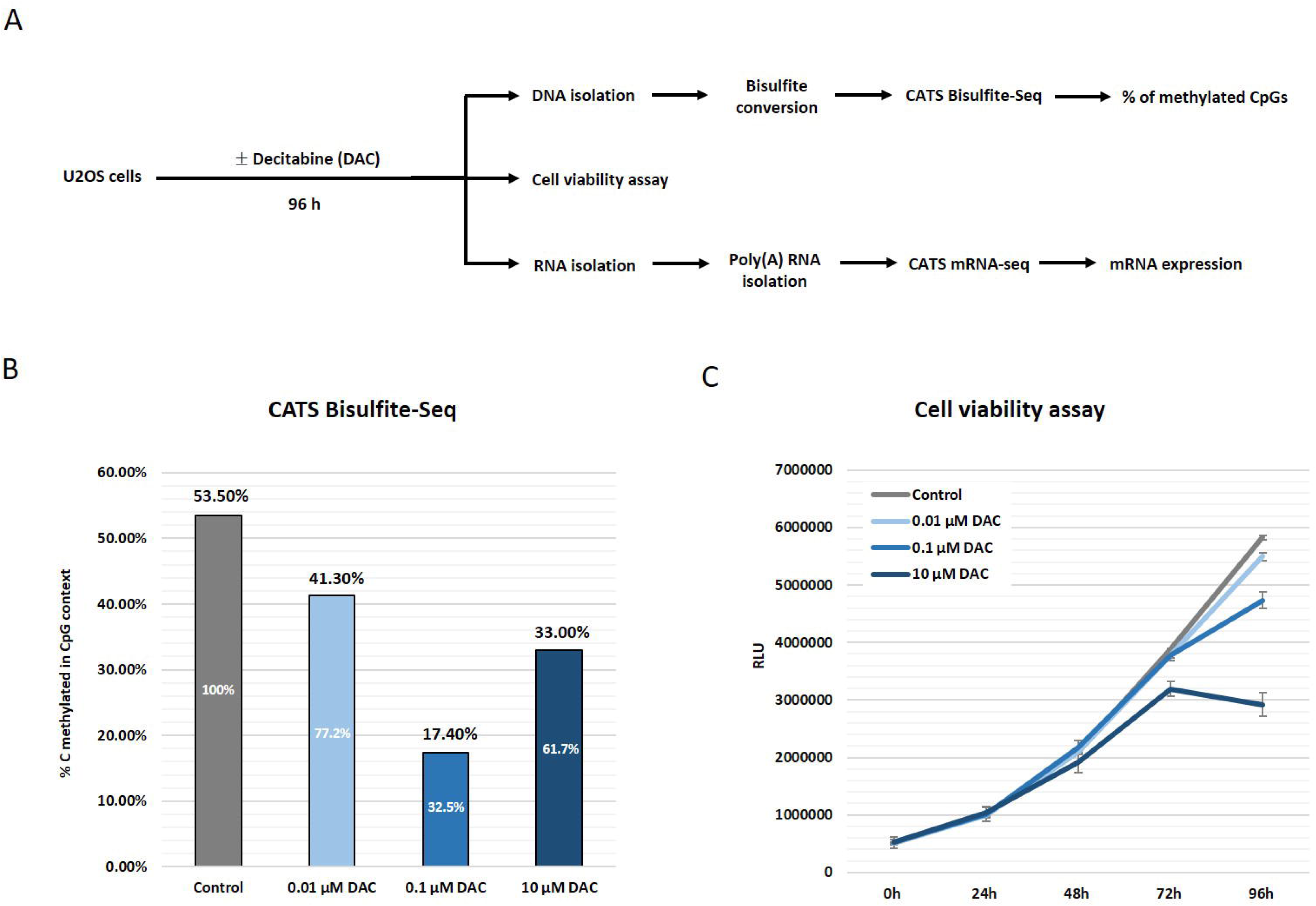
Differential impact of DAC on viability and CpG methylation status of U2OS cells. **(A)** The workflow of the experimental setup. U2OS cells in culture were treated with varying concentrations of DAC. After 96 hours DNA and RNA were isolated and used for bisulfite-DNA-seq and mRNA-seq respectively. Cell viability was measured every 24 hours after DAC treatment. **(B)** The graph demonstrating the overall percent of methylated CpG dinucleotides in control cells and cells treated with varying DAC doses. Note, low 0.1 μM DAC mediates approximately 2-fold higher hypomethylation as compared to high 10 μM doses. **(C)** Cell viability assay performed over the 96-hour course of DAC treatment of U2OS cells. Note, dose-dependent impairment of cell viability after DAC treatment.

In this study, we did not address how uniform the hypomethylation occurred over the genome of U2OS cells treated with different DAC concentrations, and merely aimed to assess the change in overall CpG methylation. Indeed, assessing demethylation efficiency over different genomic regions would require significantly higher coverage of whole-genome bisulfite sequencing. Previous data generated with methylation microarrays have suggested that the extent of hypomethylation could be non-random because the regions with a higher degree of baseline CpG methylation (e.g. gene bodies) appeared hypomethylated to the higher extent [Hagemann et al, 2012; Yan et al, 2012; Kclo et al, 2013]. On the contrary, whole-genome bisulfite deep sequencing indicated a near-uniform decrease in CpG methylation across the whole genome (while promoters, CpG islands, and 5′ UTRs underwent a slightly more modest decrease) [Lund et al, 2014]. Taken into account the above reports, we assumed that, at least, identical DNA loci to be hypomethylated with similar efficacy.

### High-throughput gene expression analysis of U2OS cells after DAC treatment

On the following step, mRNA was isolated from control and DAC treated U2OS cells, converted into NGS libraries using Capture and Amplification by Tailing and Switching (CATS) method [Turchinovich et at, 2014] and sequenced on the Illumina HiSeq platform (Supplementary Fig. 2A). The raw reads from FASTQ files have been trimmed from the remaining adapter sequences, aligned to the human reference transcriptome and counted as described in the Material and Methods section. The raw counts obtained from two biological replicates demonstrated a remarkable correlation in both controls (Supplementary Fig. 2B) and DAC treated cells (Supplementary Fig. 3). To further benchmark the accuracy of CATS mRNA sequencing, we have compared the relative abundance of four human Argonaute mRNAs (AGO1, AGO2, AGO3, and AGO4) as well as two most commonly used housekeeping genes (GAPDH and ACTB) measured by CATS mRNA-seq (Supplementary Fig. 2C) and RT-qPCR (Supplementary Fig. 2D). Both CATS mRNA sequencing and RT-qPCR experiments showed similar results regarding the relative expression of the Argonautes as well as both housekeeping genes.

**Figure 2.**
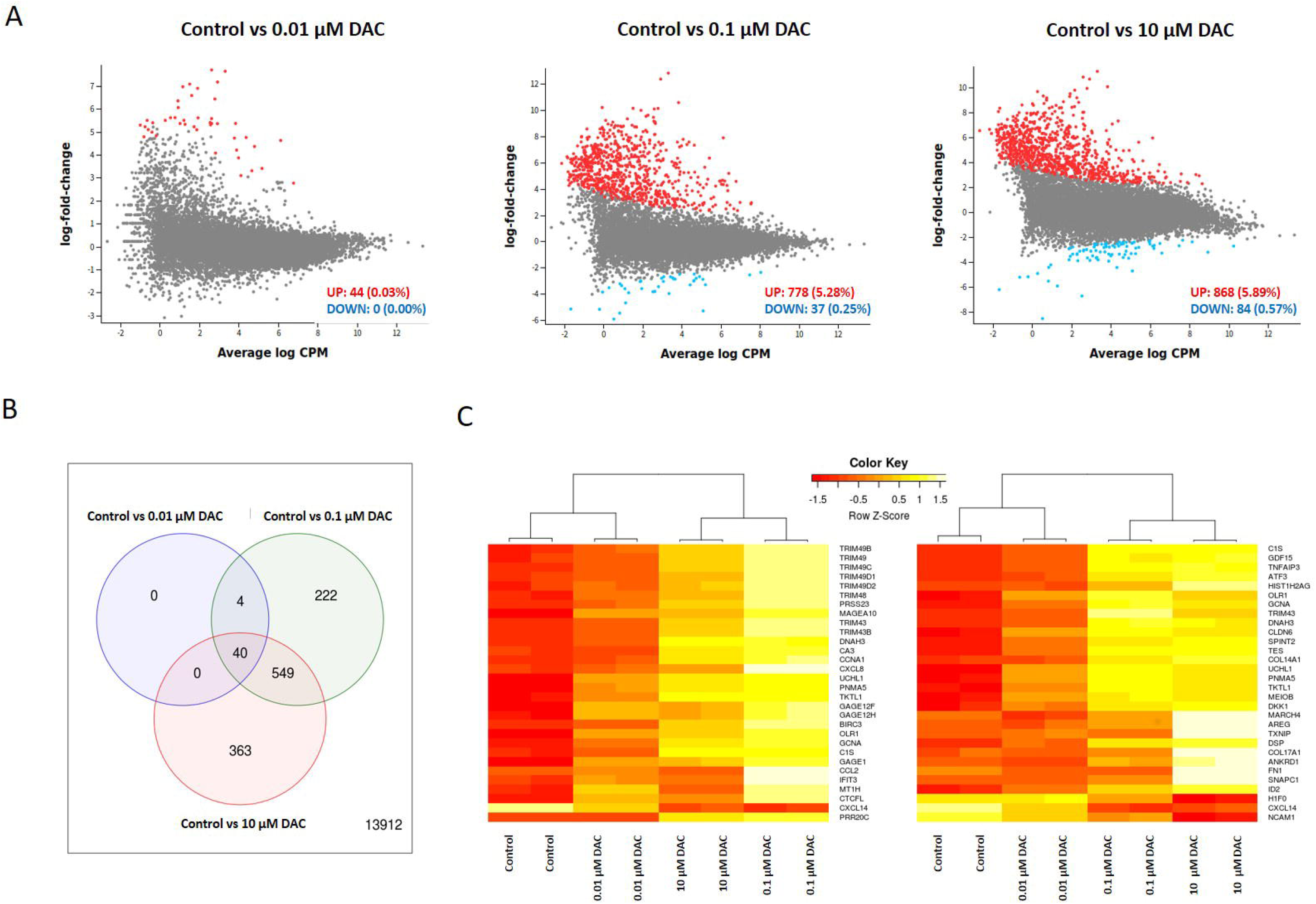
Impact of decitabine on gene expression in U2OS cells. **(A)** Mean-difference (MD) plots showing the log-fold change and the average abundance of each gene between control (untreated) cells and the cells treated with 0.01 μM, 0.1 μM and 10 μM DAC. Genes which showed statistically significant (adj-p-value < 0.05) upregulation in the treated cells (log-fold-change ≥ 2) are shown in red, while the downregulated (log-fold-change ≥ −2) genes are indicated as blue dots. Note, DAC treatment induces a very limited number of differentially expressed genes of total 15090 remaining after expression cut-off filtering (Supplementary Fig. 3). **(B)** Venn diagram showing the overlap of DE genes between cells treated with three different DAC doses. **(C)** Heatmaps of log-CPM values for top 30 DE genes (ranked by adjusted p-values) in control vs. 0.1 μM DAC (left) and control vs. 10 μM DAC (right). Expression across each gene (or row) have been scaled so that mean expression is zero and standard deviation is one; the heatmaps also rearrange the order of genes to form blocks of similar expression. Note, the heatmaps correctly cluster samples into treatment conditions and replicates.

**Figure 3.**
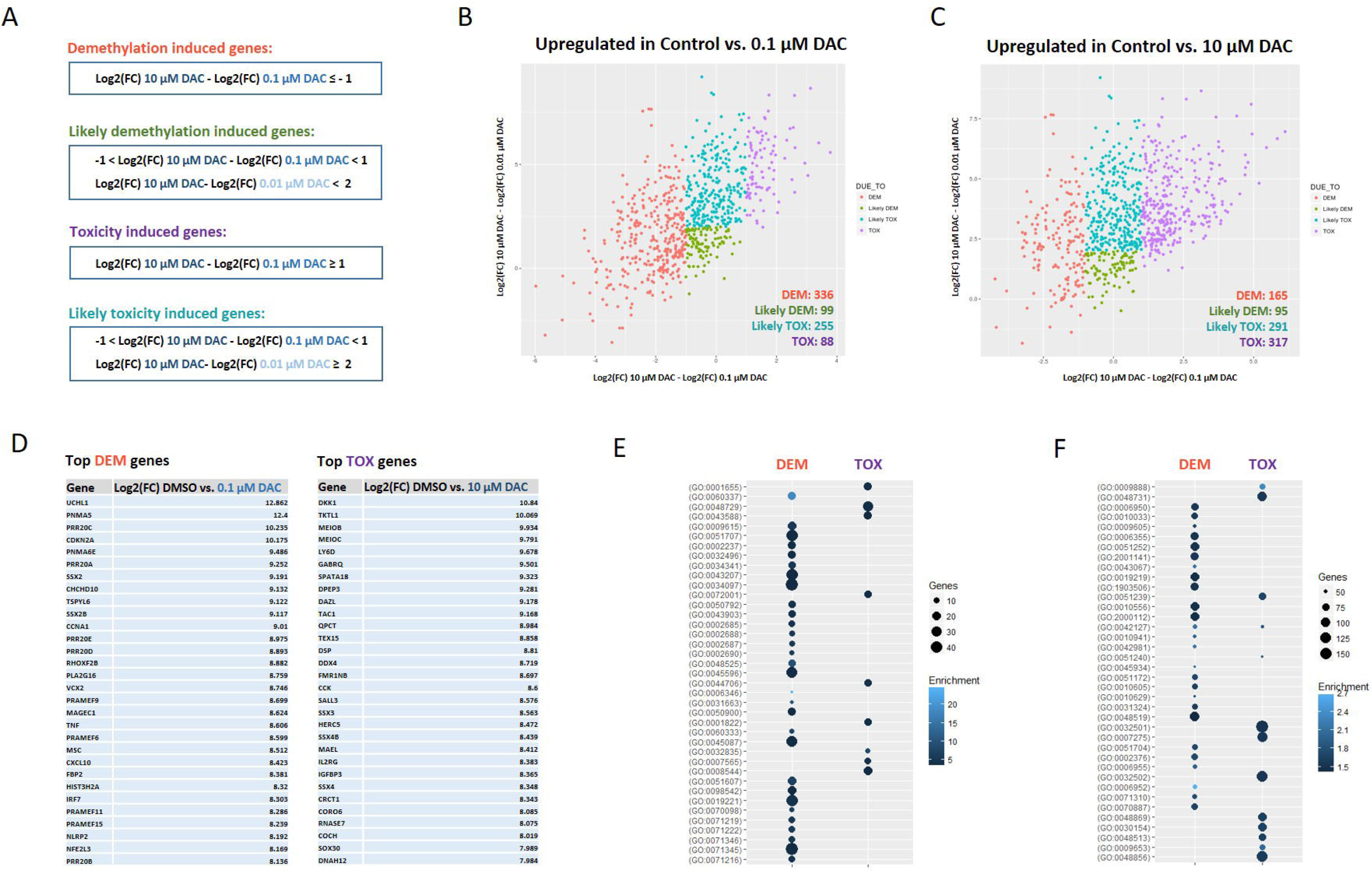
Dissection of hypomethylation and toxicity-induced mRNA expression in U2OS cells. **(A)** Criteria applied to differentiate the hypomethylation and toxicity-induced genes. **(B, C)** Scatter plots showing the distribution and the number genes upregulated due to demethylation (DEM) or “likely” demethylation (Likely DEM) as well as toxicity (TOX) or “likely” toxicity (Likely TOX) in 0.1 μM DAC and 10 μM DAC treated cells. **(D)** The lists of top 30 demethylation (left) and toxicity-induced (right) genes as ranked by log-fold-change in 0.1 μM DAC and 10 μM DAC treated cells, respectively. **(E, F)** Gene ontology analysis graphs showing the enriched GO terms for biological processes to which the identified 317 toxicity and 336 demethylation induced genes are involved.

Before performing a differential expression (DE) analysis, the filtered raw gene-level counts were transformed to log2 counts-per-million (log CPM) and normalized with the method of trimmed mean of M-values (TMM) [Robinson and Oshlack 2010] using R/Bioconductor package edgeR (Supplementary figure 4A). The R/Bioconductor Glimma package was used for the unsupervised clustering and generating an MDS plot where the distance between all RNA-seq samples can be explored (Supplementary figure 4B). The lists of differentially expressed genes between control and DAC treated U2OS cells were obtained using R/Bioconductor package edgeR and visualized as mean-difference (MD) plots using R/Bioconductor package Glimma (Fig. 2A).

**Figure 4.**
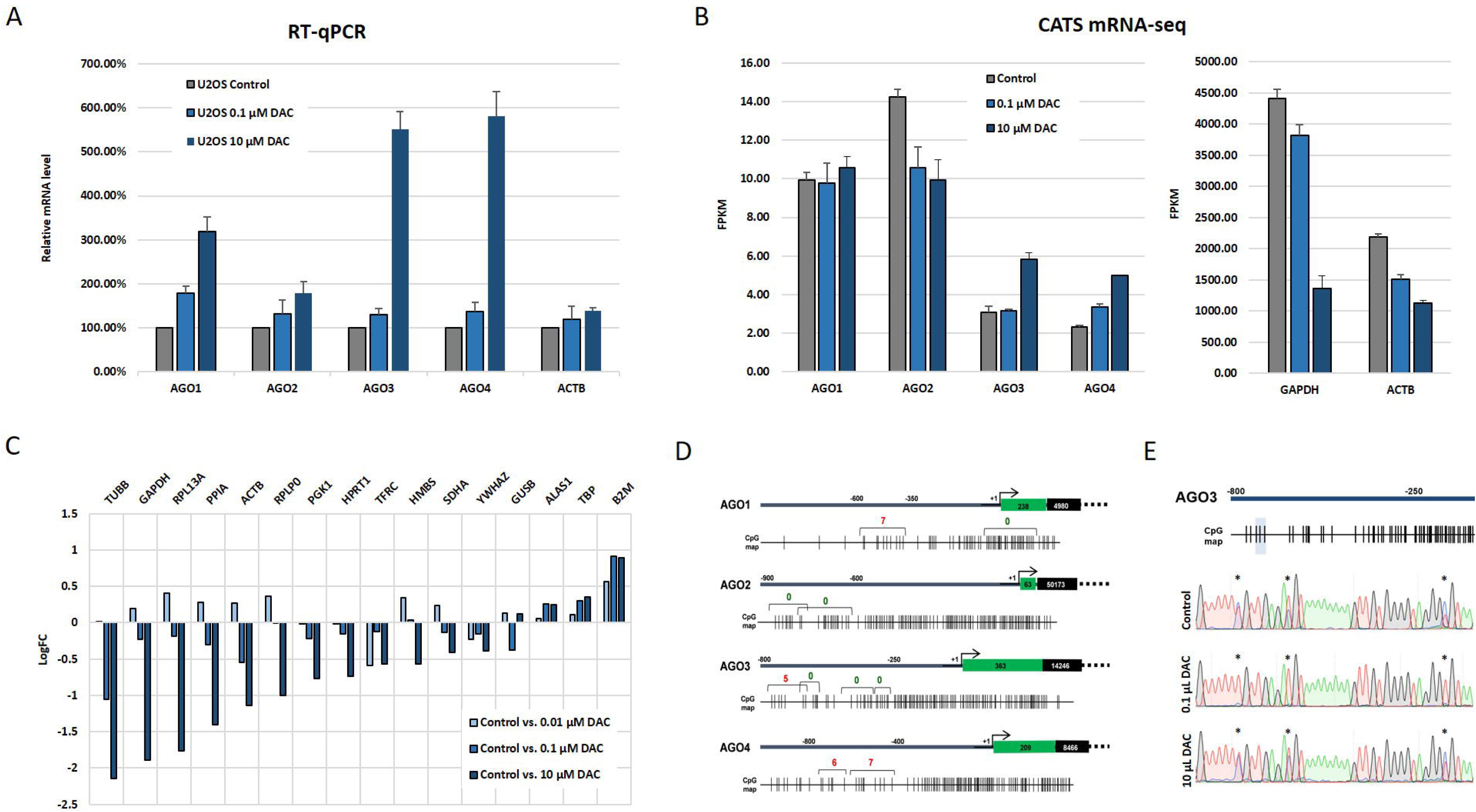
Discrepancy between RT-qPCR and RNA-seq gene expression data due to the downregulation of reference genes. **(A)** TaqMan RT-qPCR analysis of the relative abundance of human AGO1, AGO2, AGO3, AGO4 and ACTB mRNAs between control U2OS cells and cells treated with DAC for 96 hours. Ct values obtained after RT-qPCR using AGO-specific TaqMan assays were normalized on the Ct values of GAPDH mRNA and presented as relative mRNA level compared to mRNA level in control cells (taken as 100%). Each bar represents mean + S.D of 2 biological replicates. **(B)** The relative abundance of human AGO1, AGO2, AGO3, AGO4, ACTB and GAPDH mRNAs between control U2OS cells and cells treated with DAC for 96 hours measured by CATS mRNA-seq experiments. The expression values for each gene are present as FPKM (fragments per kilobase of transcripts per million of mapped reads). Each bar represents mean + S.D of 2 biological replicates. **(C)** Differential expression data of 16 common house-keeping genes between control and DAC treated U2OS cells extracted from CATS mRNA-seq experiments. The data is present as log2-fold-change of each gene in DAC treated cells vs. controls. **(D)** CpG map of the promoter regions of human AGO1, AGO2, AGO3 and AGO4 genes. The location of first exon and first intron of each Argonaute gene is shows as green and black boxes respectively with the indicated length base pairs. The brackets designate the DNA regions which were analyzed with Sanger bisulfite sequencing. The number above each bracket indicates the number of methylated CpG pairs in a given region. A CpG pair was considered “methylated” if at least 30% of the cytosines were evident in the corresponding CpG dinucleotide on a Sanger chromatogram. **(E)** Sanger chromatograms obtained after bisulfite sequencing of the region containing 3 CpG dinucleotides in the AGO3 promoter region in control and DAC treated U2OS cells. Note, significantly lower demethylation of the CpG dinucleotides in 10 μM DAC as compared to 0.1 μM DAC treated cells.

After installing the log2-fold-change (LFC) threshold for a gene to be referred as “differentially expressed (DE)” equal 2 (corresponds to 4-fold relative difference), the proportion of total DE genes appeared surprisingly limited even in cells treated with high DAC doses (with no more than 5-7% of detected genes affected) (Fig. 2A). Thus, ultra-low (0.01 μM) DAC concentrations induced expression of only 44 mRNAs, while no gene was significantly downregulated. The number of upregulated transcripts in 0.1 μM and 10 μM DAC treated cells vs. controls constituted 778 and 868 respectively; while the downregulation was detected in only 37 and 84 genes respectively (Fig. 2A). Importantly, the expression of as many as 363 genes was significantly altered in cells treated with high (10 μM) but not with low (0.1 μM) DAC doses, indicating that these changes could be due to DAC-associated toxicity rather than CpG hypomethylation (Fig. 2B). Likewise, DE of 226 transcripts was found in cells incubated with 0.01 μM and 0.1 μM DAC but not with high (10 μM) concentrations of the drug, indicating the effect to be related to hypomethylation. The observation that the expression of some transcripts was clearly altered in one treatment condition but not in the other was also evident from the heatmaps showing the hierarchical clustering based on top 30 DE genes (ranked by adjusted p-value) from the control vs. 0.1 μM DAC and control vs. 10 μM DAC contrasts (Fig. 2C). Importantly, 589 genes were significantly altered in cells treated with both low and high DAC doses (Fig. 2B), and therefore, their DE could not be unambiguously assigned to either of two modes of DAC action.

### Dissection of hypomethylation and toxicity induced genes after DAC treatment

The above DE analysis indicated that a certain number of genes in U2OS cells was upregulated due to CpG hypomethylation status, while the other transcripts could have been induced by DAC-mediated toxicity (Fig. 2B, 2C). Based on the observation that increasing DAC concentrations showed the U-shaped DNA hypomethylation efficiency (while, at the same time, maintaining a dose-dependent impact on cell viability) we have selected the criteria for dissecting genes upregulated due to CpG demethylation from genes induced by DAC toxicity (Fig. 3A). The rationale behind the selected criteria was based on an extent to which a given DAC concentration affected global DNA demethylation and cell viability (Fig. 1A, 1B). Specifically, any gene showing at least 2-fold higher upregulation in 0.1 μM DAC as compared to 10 μM DAC treated cells was considered “demethylation induced” (since the 0.1 μM DAC induced approximately 2-fold higher global DNA hypomethylation level as compared to 10 μM DAC). Likewise, a gene was considered “toxicity induced” if it showed at least 2-fold higher upregulation in 10 μM DAC as compared to 0.1 μM DAC treated samples (since the 10 μM DAC induced approximately 2-fold higher impairment of cells viability as compared to 0.1 μM DAC). Finally, the genes showing similar (below 2-fold) changes in 0.1 μM and 10 μM DAC treated cells were further split on “likely demethylation induced” and “likely toxicity induced” based on their combined behavior in cells incubated with ultra-low 0.01 μM and high 10 μM DAC concentrations. Thus, genes which demonstrated less than 4-fold difference between 10 μM and 0.01 μM DAC treated cells were assigned as “likely demethylation induced”, while the genes having more than 4-fold higher expression in 10 μM as compared to 0.01 μM DAC treated cells were assigned as “likely toxicity induced”. The rationale behind the described criteria for splitting similarly upregulated genes was based on the notion that high 10 μM DAC concentrations induced similar DNA demethylation levels (33.00% vs 41.30%) but more than four times higher impairment in cells viability as compared to cells treated with ultra-low 0.01 μM DAC dose (Fig. 1A, 1B).

In 0.1 μM DAC treated cells (that showed the highest degree of DNA hypomethylation) 336 out of 778 upregulated genes satisfied the criteria for being “demethylation induced”, while only 88 transcripts were attributed to “toxicity induced” (Fig. 3B). Among the total of 868 upregulated genes in the 10 μM DAC treated group, the number of demethylation induced genes was significantly lower (168 genes), while as many as 317 genes were activated due to cell toxicity (Fig. 3C). Interestingly, the significant proportion of genes were overexpressed to a similar extent (−1<ΔLog2(FC)<1) in both low and high DAC treated cells and were classified as either “likely demethylation induced” (99 and 95 genes in 0.1 μM and 10 μM DAC treated cells respectively) or “likely toxicity induced” (255 and 291 genes in 0.1 μM and 10 μM DAC treated cells respectively). The top 30 (based on log2-fold-change) demethylation and toxicity induced genes are listed on Figure 3D, and are known to participate in various pathways including cell proliferation, apoptosis, and immune response. Besides, subsequent GO enrichment analysis (performed using Panther [Mi et al, 2015]) clearly stratified the 336 demethylation and 317 toxicity induced transcripts in terms of their involvement in certain biological processes (Fig. 3E, 3F).

### DAC induced differential expression of a limited number of human oncogenes and tumor-suppressor genes

The analysis of all differentially expressed genes including the biological pathways in which they are involved goes well beyond the scope of this work. However, we have combined the list of most known human oncogenes and tumor-suppressors (Supplementary Table 1) and identified those which showed significant alteration in each DAC treated group (Table 1). Interestingly, out of the 48 most common oncogenes, only 14 had log2(FC) over 1.8 (approximately 3.5-fold relative change) in either 0.1 μM DAC or 10 μM DAC group. Likewise, out of the 46 most common tumor suppressor genes, only 9 had significant (log2(FC) > 1.8) changes in expression in either 0.1 μM DAC or 10 μM DAC cells. Surprisingly, differential expression of only three oncogenes (TRAF1, MYC, CCNE2) and four tumor suppressors (CDKN2A, CDKN2B, TP63, TP73) could be unambiguously attributed to CpG hypomethylation (Table 1). The CDKN2A and CDKN2B were the most overexpressed tumor suppressor genes in 0.1 μM DAC treated cells and were consistently shown to be silenced by CpG island methylation in human cancer cells [Merlo et al, 1995]. Interestingly, the TP63 gene was upregulated more than 12-fold (log2(FC) = 3,703) in 0.1 μM DAC treated cells, while its close homolog TP53 (that encodes p53 protein) showed slight downregulation (log2(FC) = - 0.619) in the same cells (Supplementary Table 1). Among human oncogenes, only TRAF1 (log2(FC) = 5.742) and MYC (log2(FC) = 1.858) were upregulated due to demethylation, while CCNE2 was downregulated (log2(FC) = −1.82) in 0.1 μM DAC treated cells. The expression of the other three oncogenes (FOSB, SOX5, and CDK15) was altered possibly due to CpG demethylation, however, only FOSB was upregulated. The remaining eight oncogenes (SOX1, SOX2, SOX10, SOX30, SYK, MDM2, NTRK2, GLI1) and five tumor suppressors (CDH1, ST14, CDKN1A, FAS, CDKN2C) showed toxicity-dependent patterns of differential expression changes (Table 1). The fact that the induction of CDKN1A gene by 5-azacitidine requires DNA damage had been also previously reported [Jiemjit et al, 2008].

**Table 1.** The list of oncogenes and tumor-suppressor genes which showed statistically significant (adj-p-value≤0.05) differential expression (log-fold-change ≥ 1.8 in any of the three treatment conditions) in U2OS cells after decitabine (DAC) treatment.

| Tumor-suppressor gene | Encoded proteins | fpkm (control) | DAC 0.01 | DAC 0.1 | DAC 10 | Mode | Function |
| --- | --- | --- | --- | --- | --- | --- | --- |
| CDKN2A | p16 and p15ARF | 0 | 7.093 | 10.175 | 7.331 | DEM | promotes cell cycle arrest |
| CDKN2B | p15INK4b | 0.37 | 2.983 | 3.879 | 2.241 | DEM | inhibits cell cycle G1 progression |
| TP63 | p63 | 0.25 | -0.083 | 3.703 | 1.893 | DEM | regulates apoptosis |
| CDH1 | E-cadherin | 0.48 | 0.374 | 2.948 | 3.348 | LikelyTOX | promotes cell-cell adhesion |
| ST14 | matriptase | 0.085 | 1.358 | 2.77 | 4.544 | TOX | degrades extracellular matrix |
| CDKN1A | p21Cip1 | 44.05 | 0.478 | 1.951 | 2.376 | LikelyTOX | promotes cell cycle arrest |
| FAS | Fas receptor | 11.1 | -0.044 | 1.24 | 3.16 | TOX | promotes apoptosis |
| TP73 | p73 | 1.85 | 0.24 | -1.814 | -0.553 | DEM | promotes apoptosis |
| CDKN2C | p18(INK4c) | 24.6 | 0.431 | -1.974 | -2.023 | TOX | controls cell cycle G1 progression, suppress cell growth |
| Oncogene | Encoded proteins | fpkm (control) | DAC 0.01 | DAC 0.1 | DAC 10 | Mode | Function |
| SOX30 | Sox30 | 0 | 1.403 | 6.497 | 7.989 | TOX | determination of cell fate |
| TRAF1 | NF receptor-associated factor 1 | 0.08 | 1.91 | 5.742 | 4.137 | DEM | activates of MAPK8/JNK and NF-kappaB |
| SOX10 | Sox10 | 0 | 1.035 | 4.283 | 6.126 | TOX | determination of cell fate |
| SOX1 | Sox1 | 0.06 | -1.922 | 4.165 | 3.733 | TOX | determination of cell fate |
| FOSB* | FosB | 0.05 | 1.859 | 3.232 | 3.115 | LikelyDEM | interacts with proteins of JUN family |
| MYC | c-Myc | 29.65 | 0.281 | 1.858 | 0.267 | DEM | promotes cell transformation |
| SYK | Tyrosine-protein kinase SYK | 0.13 | 0.159 | 1.655 | 3.333 | TOX | transmits signals from a variety of cell surface receptors |
| MDM2 | Mdm2 | 10.95 | -0.172 | 1.027 | 2.294 | TOX | negative regulator of the p53 |
| NTRK2 | TrkB | 0.06 | -1.103 | 0.965 | 2.207 | TOX | cell differentiation |
| GLI1 | Glioma-associated oncogene | 4.19 | -0.458 | -1.212 | -2.218 | TOX | determination of cell fate |
| CCNE2 | Cyclin E2 | 7.08 | -0.584 | -1.82 | -0.772 | DEM | regulates G1/S transition |
| SOX5 | Sox5 | 1.31 | -1.279 | -2.283 | -2.769 | LikelyDEM | determination of the cell fate |
| SOX2 | Sox2 | 10.13 | -0.439 | -2.673 | -3.502 | TOX | maintaining self-renewal and pluripotency |
| CDK15 | Cyclin Dependent Kinase 15 | 8.6 | -1.8 | -3.431 | -3.673 | LikelyDEM | antiapoptotic activity |

### Most housekeeping genes used for RT-qPCR are downregulated after DAC treatment

In an attempt to validate the accuracy of CATS mRNA-seq protocol employed in the current experimental setup, we have measured the relative expression of four human Argonaute transcripts (AGO1, AGO2, AGO3, AGO4) as well as ACTB genes in controls and DAC treated U2OS cells with RT-qPCR (Fig. 4A). The common house-keeping GAPDH gene was used as a reference to normalize the expression values obtained in RT-qPCR. However, a significant discrepancy between CATS mRNA-seq and RT-qPCR data have been observed (Fig. 4B). Thus, treatment of U2OS cells with 10 μM DAC significantly increased the expression of endogenous AGO1, AGO3 and AGO4 mRNAs (relative to the mRNA of house-keeping GAPDH gene), while the increase in AGO2 mRNA was less profound (Fig. 4A). However, no elevation of AGO1 mRNA content was observed in the same RNA samples measured by CATS mRNA-seq, and the expression of AGO2 was even slightly downregulated (Fig.4B). Finally, AGO3 and AGO4 transcripts showed significantly modest upregulation in CATS mRNA-seq experiments as compared to RT-qPCR, while both ACTB and GAPDH were downregulated 2-3-fold. Importantly, the observed upregulation of Argonatues and the downregulation of house-keeping GAPDH and ACTB genes was toxicity induced, since their changes in low (0.1 μM) DAC does treated cells were much less profound (Fig. 4A, 4B).

Along with GAPDH and ACTB, several other house-keeping genes, which are widely used for qPCR normalization, including TUBB, RPL13A and PPIA were downregulated more than 2-fold in U2OS cells treated with 10 μM DAC (Fig. 4C). The TUBB transcript showed the highest downregulation rate among all analyzed house-keeping genes (log2(FC) = −1.055 and log2(FC) = −2.148 in 0.1 μM DAC and 10 μM DAC treated cells, respectively), while the BM2 gene showed on the contrary almost 2-fold upregulation after treatment with both 0.1 μM and 10 μM DAC doses (log2(FC) = 0.913 and log2(FC) = 0.894 respectively) (Fig. 4C). The other analyzed house-keeping genes including SDHA, YWHAZ, GUSB, ALAS1 and TBP were affected to a much lower extent upon DAC treatment (log2(FC) between -0.5 and 0.5), and could, therefore, serve as more reliable candidates for qPCR normalization in similar experimental setups (Fig. 4C).

Both the U-shaped curve of DAC demethylation efficacy and its toxicity-related impact on the expression of common house-keeping genes strongly suggest that studies involving DAC treatment can lead to erroneous conclusions regarding epigenetic silencing of a given locus. For instance, CpG demethylation could have been erroneously considered a direct cause of Argonaute genes upregulation, since RT-qPCR data demonstrated their strong and dose-dependent induction after treatment with the increasing DAC concentrations (Fig. 4A). Furthermore, AGO1, AGO3 and AGO4 genes but not AGO2 gene contained multiple methylated CpG dinucleotides 600-900 bp upstream of their TSSs (Fig. 4D) hinting on the putative epigenetic silencing of at least AGO1, AGO3 and AGO4 transcripts. In fact, upregulation Argonautes after DAC treatment could not be a consequence of DNA hypomethylation, but rather attributed to the toxicity of the agent. The latter also became evident after the analysis of DNA demethylation efficacy of different DAC doses both on the global DNA level (Fig. 1B) and locally on AGO3 gene promoter (Fig. 4E).

## Discussion

The role of CpG DNA methylation in gene expression has been the subject of intense investigations over the last decades. The majority of DNA methylation in mammalian genomes occurs at cytosine residues which are followed by guanines (CpG sites). Furthermore, multiple genomic regions harbor the so-called “CpG islands” – the regions containing CpG sites in high densities. While most CpG pairs in mammalian genomes are methylated, approximately half of human promoters contain CpG islands which methylation density negatively correlates with gene expression [Takai and Jones, 2002; Taby and Issa 2010; Jones, 2012; Smith and Meissner 2013]. It was originally assumed low methylated promoters are permissive for gene expression. However, later reports have suggested that lower DNA methylation at gene promoters could be rather a consequence of the robust gene expression from a given promoter [Flotho et al, 2009; Jones, 2012]. Indeed, active promoters could be resistant to DNA methylation both de-novo and post-replicative due to shielding by the associated RNA polymerases, transcription factors, and chromatin remodeling proteins. At the same time, the observed DNA hypermethylation of transcription start sites and other regulatory loci of the silenced genes could be due to the higher accessibility of such regions for DNA methyltransferases [Fahy et al, 2012].

The cytosine analog decitabine (DAC, 5-azadCyd) remains the most widely used chemical for experimental reduction of CpG methylation levels [Jones and Taylor, 1980]. While early studies have shown the correlation of DAC-induced DNA demethylation and activation of silent genes, multiple recent high-throughput gene expression analyses have failed to confirm a strong association between changes in methylation status and gene expression [Klco et al, 2013; Lund et al, 2015; Flotho et al, 2009; Tsai et al, 2012; Mossman et al, 2010; Schmelz et al, 2005; Weber et al, 2007; Claus et al, 2013; Ramos et al, 2015; Tobiasson et al, 2017]. Furthermore, DAC treatment has been shown to affect the expression of only a small subset of protein-coding genes despite a robust genome-wide hypomethylation [Klco et al, 2013; Ramos et al, 2015]. At the same time, recently generated high-resolution genomic maps of DNA methylation confirmed that most DNA regulatory sequences are indeed unmethylated when active [Schübeler 2015]. The one explanation for this could be the fact that removal of 5-methylcytosine residues may not be sufficient for a gene reactivation due to the presence of other modifications including histone modifications and chromatin remodeling [Si et al, 2010].

Besides CpG demethylation, DAC incorporation into the DNA is highly cytotoxic due to the covalent and irreversible binding of DNMT1 that results in DNA damage [Flatau et al, 1984; Davidson et al, 1992; Jutterman et al, 1994; Jiemjit et al, 2008]. Therefore, the toxicity impact of DAC has to be taken into account to evade the erroneous conclusions regarding the epigenetic silencing of genes. Importantly, due to the dual-mode of DAC action on living cells, a simple correlation of a promoter DNA demethylation with a gene expression changes cannot be used as proof that a given gene is silenced by CpG methylation. Furthermore, to our knowledge, the relation between DAC-induced DNA demethylation and cytotoxicity remained largely unresolved in the previous experiments. Therefore, it remained important to establish a model that allows dissecting the genes which expression is altered by DNA hypomethylation from cytotoxicity related to DAC-mediated DNA damage.

Due to the limited availability of suitable tools for genome-wide methylation analysis, the studies of DNA hypomethylation by DAC were initially restricted to the analysis of selected DNA regions in various reports [Mund et al, 2005; Yang et al, 2006]. In this work, we performed genome-wide methylation profiling of cultured U2OS cells using recently described CATS bisulfite DNA-seq protocol [Turchinovich et al, 2014]. In contrast to other techniques, the CATS method allows using bisulfite-converted single-stranded DNA as input directly, has little (if any) bias to particular genomic regions and requires only a limited amount of DNA input [Turchinovich et al, 2014]. Although we did not aim to investigate global methylation status at a single-base resolution, CATS bisulfite DNA-seq confirmed the U-shaped curve of the de-methylation efficacy of the increasing DAC concentrations, a phenomenon which was previously demonstrated only for LINE DNA region in a panel of cancer cell lines [Qin et al, 2007]. Specifically, we show that low 0.1 μM DAC induced significantly stronger DNA hypomethylation as compared to ultra-low 0.01 μM and high 10 μM drug doses, while the impact of increasing DAC concentrations on cell viability was linear. We have further utilized CATS mRNA-seq protocol to explore the effects of the above DAC doses on gene expression in U2OS cells. Besides, by analyzing the expression of selected reference mRNAs, we confirmed that CATS mRNA-seq allows estimation of relative gene expression on a high-throughput level with accuracy similar to RT-qPCR (Supplementary Fig.2). Finally, the U-shaped behaviors of DAC response allowed putative dissection of gene expression changes occurring specifically due to hypomethylation and cell toxicity.

Although we did not address the molecular mechanisms responsible for lower hypomethylation impact of high DAC doses, the latter could have induced much cell-cycle arrest much earlier and render stronger inhibition of DNA replication, which in turn manifested in lower CpG hypomethylation. At the same time, low DAC doses might have not induced significant cell-cycle arrest while was still able to maintain efficient trapping of DNA methyltransferases. Consequently, our mRNA-seq results have shown that a higher proportion of genes were upregulated due to epigenetic mechanisms in cells treated with low DAC doses, while the cytotoxic gene expression responses prevailed at high doses of the drug. Specifically, the number of demethylation-induced genes in cells treated with high decitabine concentrations was approximately 50% lower as compared to cytotoxicity-induced genes, while at low DAC doses the number of demethylation-induced genes was 1.5-fold higher. In accordance with the previous reports, DAC also induced downregulation of some genes, however, their number was significantly lower [Gius et al, 2004; Schuebel et al, 2007].

Interestingly, DAC treatment of U2OS cells induced expression changes in a moderate number of transcripts (between 4.1%-4.8%, of total 19835 genes in Ensemble reference database). Besides, upregulation (or restoration) of mRNA expression of more than half of these genes could not be directly attributed to DNA methylation changes but rather to cell toxicity. The limited transcriptional consequences of DNA demethylation has been previously reported by other studies which mainly used microarrays platforms [Karpf et al, 2004; Shi et al, 2003; Liang et al, 2002; Al-Romaih et al, 2007; Kim et al, 2006; Lund et al, 2014; Klco et al, 2013; Ramos et al, 2015].

Previous studies have also indicated that hypermethylation of CpG islands and CpG shores can be responsible for silencing certain tumor suppressor genes in many human cancers [Jones and Baylin 2007; Baylin and Jones, 2011; Irizarry et al, 2009; Plass et al, 2013; Taby and Issa 2010; Jelinek et al. 2012]. Therefore, the anti-tumor impact of DAC was frequently attributed to the demethylation of regulatory regions of silenced tumor-suppressors and consequent cell growth arrest, apoptosis and/or differentiation [Wilson et al, 1983; Jones et al, 1983; Momparler et al, 1985; Willemze et al, 1993]. For instance, CDKN2B and CDH1 (E-cadherin) genes have been reported to be silenced by CpG methylation in cancer cells [Paul et al, 2010; Graff et al, 1995; Yoshiura et al, 1995; Nam et al, 2004]. While our data confirm the demethylation-dependent upregulation of CDKN2B tumor suppressor in U2OS cells, the reactivation of the CDH1 gene was clearly due to DAC-mediated cytotoxicity. Furthermore, we found that another tumor-suppressor gene CDKN1A was upregulated several-fold by toxicity-related mechanism, consistent with previous reports showing that CDKN1A induction requires DNA damage [Jiemjit et al, 2008]. Surprisingly, while in our experimental setup, DAC significantly affected the expression of 23 different oncogenes and tumor suppressors, only 3 tumor suppressor genes (CDKN2A, CDKN2B and TP63) and 2 oncogenes (TRAF1 and MYC) were activated due to DNA demethylation in U2OS cells (Table 1, Supplementary Table 1). Another research group had also reported a marked induction of the CDKN2A gene in U2OS cells upon transient DAC treatment that was correlated with the level of demethylation over a defined region of DNA on the CDKN2A promoter [Badal et al, 2008]. In our work, CDKN2A was one of the most strongly upregulated genes due to demethylation and was not expressed in the control cells. Furthermore, in accordance with Badal et al, we did not observe an induction of p53 gene in U2OS cells upon DAC treatment.

Overexpression of 13 different genes in U2OS cells after DAC treatment has been previously demonstrated by a microarray platform [Al-Romaih et al, 2007]. Furthermore, hypomethylation of promoters of GADD45A, PAWR, HSPA9B and PDCD5 genes have been confirmed by pyrosequencing [Al-Romaih et al, 2008]. However, our mRNA-seq experiments have confirmed the significant upregulation of only 8 out of those 13 genes. Furthermore, the upregulation of only NFKBIA and RAC2 genes was demethylation dependent, while the overexpression of the other 6 transcripts (including IGFBP6, RIPOR1 FAM65A, TNFAIP3, GADD45A, PAWR and PSG5) strongly correlated with cell toxicity response. Although we did not find the differences in expression of GAGE4 gene (which was also reported by [Al-Romaih et al, 2005]), multiple other GAGE genes (all having a high degree of sequence identity) including GAGE1, GAGE2 (A, E), GAGE10, GAGE12 (B-J) and GAGE13 were markedly overexpressed due to DNA demethylation in U2OS cells. While the exact function of GAGE proteins is presently unknown, their sequences contain the antigenic peptides which are recognized by cytotoxic T-cells and, therefore, could be related to the anti-cancer immune response.

Previous studies on multiple cancer cell lines have shown that the dominant effect of transient DAC treatment resulted in activation of various immune-related genes involved in interferon/cytokine signaling, antigen presentation and inflammation (Karpf et al. 1999; Wrangle et al. 2013; Odunsi et al. 2014; Heninger et al. 2015; Li et al. 2014; Liang et al, 2002). Interestingly, most of those genes did not have CpG island promoters and the epigenetic mechanism controlling their re-expression was not evident [Li et al, 2014]. Our data confirmed a strong induction of genes involved in the immune-response pathways following DAC treatment of U2OS cells. Specifically, multiple genes were strongly upregulated due to DNA hypomethylation including tumor necrosis factor TNF, interferon-induced GTP-binding protein MX1, various human leukocyte antigens (HLA-A, HLA-B, HLA-C, HLA-DRB3), interleukins (IL16, IL18), C-C Motif Chemokine Ligands (CCL2, CCL20, CCL3, CCL5), clusters of differentiation (ICAM1, ICAM2, CD69, CD24), interferon regulatory factors (IRF7, IRF9), interferon alpha-inducible proteins (IFI27, IFI35, IFI44, IFI44L, IFI6), interferon induced transmembrane protein IFITM1, interferon-induced helicase C domain-containing protein IFIH1, interferon-induced proteins with tetratricopeptide repeats (IFIT1, IFIT2, IFIT3, IFIT5), C-X-C motif chemokines (CXCL10, CXCL11, CXCL5, CXCL8), 2’-5’-oligoadenylate synthetases (OAS1, OAS2, OAS3, OASL), interferon-stimulated genes (ISG15, ISG20), signal transducer and activator of transcription STAT1, serum amyloid A1 (SAA1), XIAP-associated factor 1 other gene (XAF1), DExD/H-Box Helicase 58 (DDX58), TNF alpha induced protein 2 (TNFAIP2), IL15RA interleukin 15 receptor subunit alpha (IL15RA), C-C motif chemokine receptor 4 (CCR4) and NFKB inhibitor alpha (NFKBIA). However, some immune-related genes such as CD44, CCL28, ICAM4, ICAM5, GBP1, CXCL16, CXCL17, CD68, HERC5, TNFAIP3, IL6R, IL11, IL12A, IL12RB2, IL13RA2, IL1A and IL1R2 showed toxicity-specific upregulation pattern. Our findings, therefore, confirm the epigenetic silencing of the immune response-related genes by DAC in cancer cells and stresses a putative immunomodulatory role for DNA demethylation drugs in cancers.

To conclude, our study introduces an in vitro model for dissecting cytotoxic and hypomethylating impacts of DAC on gene expression in cultured cancer cells. The U-shaped curve of DAC-induced CpG hypomethylation was confirmed to occur on a global genome level, and stress the importance of using the optimal doses of DAC to maximize its epigenetic effects both for clinical and research applications. We also found that expression of multiple house-keeping transcripts was significantly affected in U2OS cells after treatment with certain DAC doses. As a result, even non-affected genes could manifest themselves as “up-regulated” on RT-qPCR experiments when their expression is normalized on a downregulated house-keeping gene. Finally, our results demonstrate the importance of accounting for DAC toxicity-related effects before making a conclusion regarding the epigenetic regulation of a gene expression.

## Materials and Methods

### Cell culture and DAC treatment

The U2OS cells were obtained from the American Type Culture Collection (ATCC) and additionally genetically authenticated by DKFZ (Heidelberg) core facility. U2OS cells were grown in DMEM supplemented with 10% FBS, 1 % non-essential amino acid and penicillin/streptomycin at 37°C in 5% CO_2_. For DAC treatment, the cells were seeded in 6-well plates at a density of 5×10^4^ cells per well respectively. Decitabine (DAC, 5′-Aza-2’-deoxycytidine) was purchased from Sigma-Aldrich, diluted in DMSO and added 12 hours after plating at 0.01 μM, 0.01 μM or 10 μM concentrations. The DMSO corresponding to its concentration in the 10 μM DAC wells was added into the control cells. The media containing freshly diluted DAC was renewed every 24 hours. Cells were harvested after 96 hours of incubation and used for RNA and DNA extraction.

### Cell viability assay

U2OS cells were seeded in opaque-walled 96-well plates and incubated for 12 hours prior to DAC addition. Control wells containing cell culture medium without cells were used to obtain a value for background luminescence. The number of viable cells in each well was determined with CellTiter-Glo^®^ Luminescent Cell Viability Assay (Promega) according to the manufacturer’s recommendations. The luminescence was read on a Tecan Microplate Reader.

### Whole-Genome Bisulfite Sequencing

Genomic DNA was isolated from control and DAC treated U2OS cells growing on the 6-well plates using QIAamp DNA Mini Kit (Qiagen), and bisulfite-converted using MethylEdge™ Bisulfite Conversion System (Promega) according to the manufacturers’ protocols. Approximately 10 ng of bisulfite-treated single-stranded DNA was directly used for NGS libraries preparation with CATS DNA-seq Kit v1.0 (Diagenode, Liege, Belgium) according to the manufacturer’s instructions. The CATS DNA libraries were multiplexed, diluted and loaded onto the HiSeq 2000 System for sequencing using 50 bp singe-end reads. Approximately 30 million raw reads were obtained per each sample. Methylation data analysis was performed using Bismark software v0.19.0 [Krueger et al, 2011]. Briefly, adapter sequences in the raw FASTQ files were trimmed with cutadapt software v1.15 as described in Supplementary Material 6. A bisulfite-converted Ensemble GRCh38 reference genome file was generated using Bismark/Bowtie2, and the CATS library sequence data were aligned to the reference genome as described in Supplementary Material 6. Methylation information was extracted from the output bam file with Bismark as described in Supplementary Material 6, and contained the percentage of methylated cytosines in CpG context observed in each sample.

### High-Throughput mRNA sequencing

Total RNA was isolated from control and DAC treated U2OS cells growing on the 6-well plates using miRNAeasy kit (Qiagen) according to the manufacturer’s protocol and eluted in 50 μL RNAse-free water. Approximately 500 ng of total RNA was used for NGS libraries preparation with CATS mRNA-seq kit (Diagenode, Liege, Belgium) according to the manufacturer’s recommendations. The CATS mRNA libraries were multiplexed, diluted and loaded onto the HiSeq 2000 System for sequencing using 50 bp singe-end reads. Approximately 20 million of raw reads was obtained per each sample and adapter sequences in the raw FASTQ files were trimmed with cutadapt software v1.15 as described in the Supplementary Material 6. The trimmed raw reads longer than 20 nt were aligned to the manually curated human Ensemble GRCh38.89 cDNA protein-coding reference transcriptome using bowtie2 mapper as described in Supplementary Material 6. The numbers of reads mapped to each transcript were extracted using eXpress software v1.5.1 [Roberts and Pachter, 2012] as described in Supplementary Material 6. The differential gene expression was quantified either manually (Supplementary Material 6) or using R/Bioconductor packages edgeR and glimma as described in [Law et al, 2016].

### RT-qPCR

Total RNA was isolated with miRNAeasy kit (Qiagen) according to the manufacturer’s protocol and eluted in 50 μL RNAse-free water. High-Capacity cDNA Reverse Transcription Kit (Applied Biosystems) was used for cDNA preparation according to the manufacturer’s protocol. Briefly, 500 ng of total RNA diluted in 10 μL of RNAse-free water was combined with 10 μL RT mastermix containing: 2 μL 10X RT Buffer, 0.8 μL 100 mM dNTPs, 10X random primers, 1 μL RNase Inhibitor, 3.2 μL Nuclease-free water and 1 μL MultiScribe™ Reverse Transcriptase. The RT reaction was incubated at 25°C for 10 min, 37°C for 120 min and 85°C for 5 min. The cDNA product was further diluted six times with RNAse-free water and used directly for Real-time qPCR. The levels of human AGO1, AGO2, AGO3, AGO4, ACTB and GAPDH mRNAs were measured using individual TaqMan Gene Expression Assays from Applied Biosystems (TaqMan Assay IDs: Hs01084653_m1, Hs00293044_m1, Hs00227461_m1, Hs01059731_m1, Hs99999903_m1 and Hs99999905_m1 respectively). Real-time PCR was performed in a 10 μL volume and included 2 μl of diluted cDNA product, 5 μl TaqMan Universal PCR Master Mix (Applied Biosystems), 0.5 μl of corresponding TaqMan Gene Expression Assay primers and 2.5 μl of nuclease-free water. Real-time qPCR was performed using the LightCycler 480 Real-Time PCR System (Roche). The reactions were incubated in duplicates in 384-well plates at 95°C for 10 min, followed by 50 cycles of 95°C for 15 sec and 60°C for 1 min. The data were analyzed with the LightCycler 480 software (Roche), determining the threshold cycle (Ct) by the second derivative max method. The efficiencies of all TaqMan Assays were estimated from standard dilution curves. The quantities of AGO2, AGO3 and AGO4 amplicons were estimated relative to AGO1 gene using the formula X _AGO1_^Ct AGO1^/X_y_^Cty^, where X_y_= amplification efficacy for the amplicons of gene y and X_AGO1_ = amplification efficacy for the amplicons of AGO1 gene.

## Authors’ contributions

AT designed the study, carried out all experiments, performed the data analysis from NGS experiments and wrote this manuscript; HMS participated in the analysis of the NGS experiments and revised the manuscript; AGT participated in the data interpretation and revised this manuscript; BB coordinated the work, participated in study design and data analysis and revised this manuscript.

## Supplementary Figure legends

**Supplementary figure 1. (A)** The graphs showing per base sequence quality scores and sequence content across all bases in the adapter-trimmed reads obtained after CATS bisulfite sequencing of DNA isolated from corresponding U2OS cells. The graphs were generated by FASTQC software. (B) A table showing the percentage of PCR duplicates in each sample.

**Supplementary figure 2. (A)** The experimental workflows of both CATS mRNA-seq and RT-qPCR experiments with the indicated timing of each step. 500 ng of total RNA was used as an input for mRNA isolation and subsequent NGS library preparation as well as for RT-qPCR; this is approximately the amount total RNA in several microliters of eluate obtainable from a confluent cells monolayer growing in a single well of a standard 24-well plate. Approximately 20 million raw reads per sample was obtained after high-throughput sequencing on Illumina HiSeq 2000 platform and demultiplexing. The raw reads were trimmed from the adapters, mapped to the human reference transcriptome and the raw count were obtained as described in Material and Methods section. Please note, both the requirement of starting material and the time requirement for the described RNA-seq and RT-qPCR are similar. **(B)** A scatter plot showing correlation of raw counts (in a log-scale) obtained from two biological replicates of CATS mRNA-seq from DMSO treated (control) cells. **(C)** Fragments per kilobase of transcript per million reads (FPKM) metrics was used to calculate the relative mRNA abundance of human AGO1, AGO2, AGO3, AGO4, GAPDH and ACTB genes in control replicates from CATS RNA-seq data. The data is presented as mean FPKM values + SD from two biological replicates. **(D)** The same RNA samples were subjected to RT-qPCR, and the relative expression of human AGO1, AGO2, AGO3, AGO4, GAPDH and ACTB genes was determined after proper accounting for amplification efficiencies. The data is presented as mean Ct values + SD from two biological replicates. Note, the relative abundances of the six transcripts measured by RT-qPCR were similar to their relative amounts measured by CATS mRNA-seq.

**Supplementary figure 3**. Scatter plots showing correlation of read counts in a log2-scale obtained from two biological replicates of CATS mRNA-seq from control and DAC treated U2OS cells. Before processing, the raw counts in each sample were first filtered from low expressed genes (cut-off: S[raw reads over all samples] > 80). Secondly, the one outlying gene (red circle) was removed from the data.

**Supplementary figure 4.** The RNA-seq counts were converted to log2-counts-per-million (log-CPM) values using the cpm function in edgeR (log-transformations used a prior count of 0.25 to avoid taking the log of zero). The normalisation was performed by the method of trimmed mean of M-values (TMM) using the calcNormFactors function in edgeR. **(A)** Boxplots of log-CPM values showing expression distributions for unnormalised data (left) and normalised data (right) for each CATS RNA-seq sample. **(B)** MDS plot of log-CPM values over dimensions 1 and 2 with samples coloured by sample groups generated by glMDSPlot function in Glimma. The distances on the plot correspond to the leading fold-change, which is the average (root-mean-square) log2-fold-change for the 500 genes most divergent between each pair of samples by default.

**Supplementary figure 5. (A)** The formula used to calculate fragment per kilobase of transcript per million reads (FPKM) values and the log2-fold-change (LFC) of differentially expressed genes. FPKM method calculations used a prior count of 1 to avoid taking the log of zero. **(B)** The graph showing correlation of differential expression of 16 housekeeping genes in control vs. DAC treated cells (in log-fold-change) calculated by manual “FPKM method” and edgeR package.

**Supplementary figure 6.** The list of Bash/Shell commands executable under Linux environment used for NGS data processing and analysis.

