## Supplementary Figures for "Dissection of demethylation and toxicity induced gene expression changes after decitabine treatment"

### Supplementary Figure 1

A

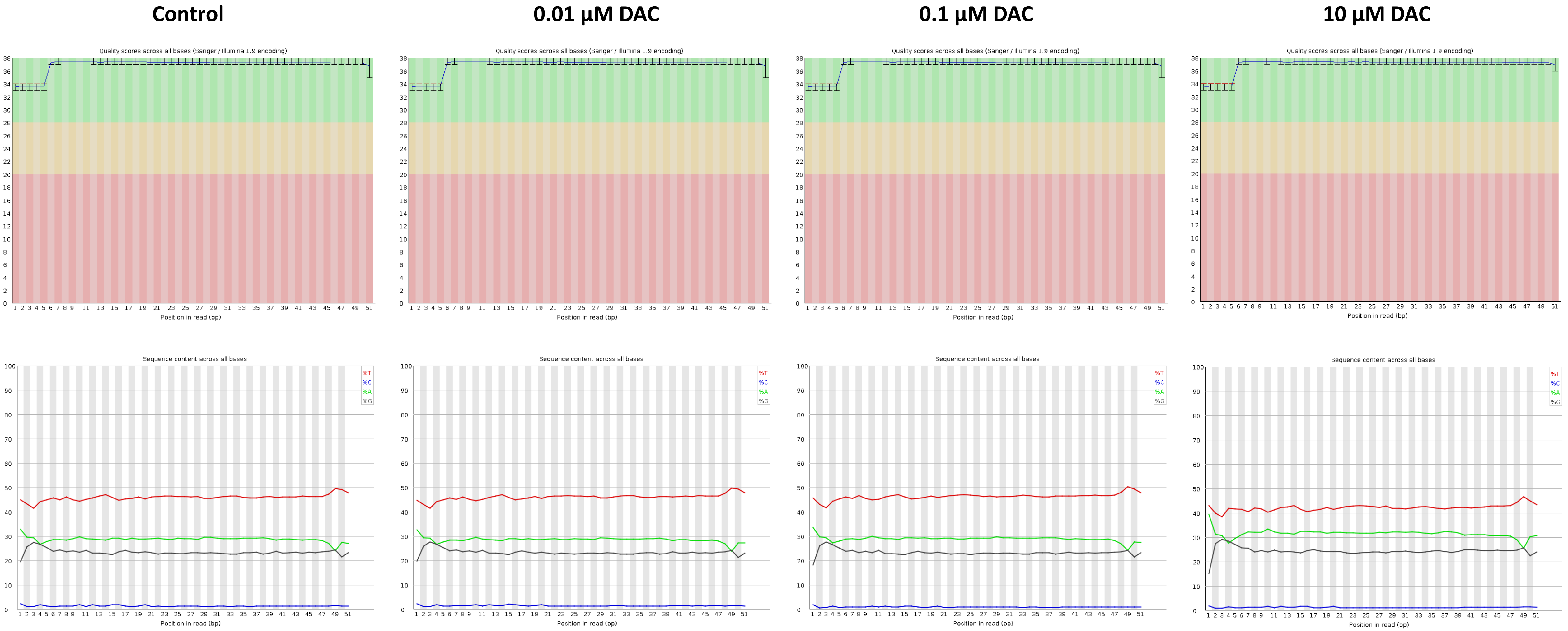


B

| | Control | 0.01 µM DAC | 0.1 µM DAC | 10 µM DAC |
|---|---|---|---|---|
| % duplicates | 6.43% | 6.39% | 6.41% | 3.32% |

### Supplementary Figure 2

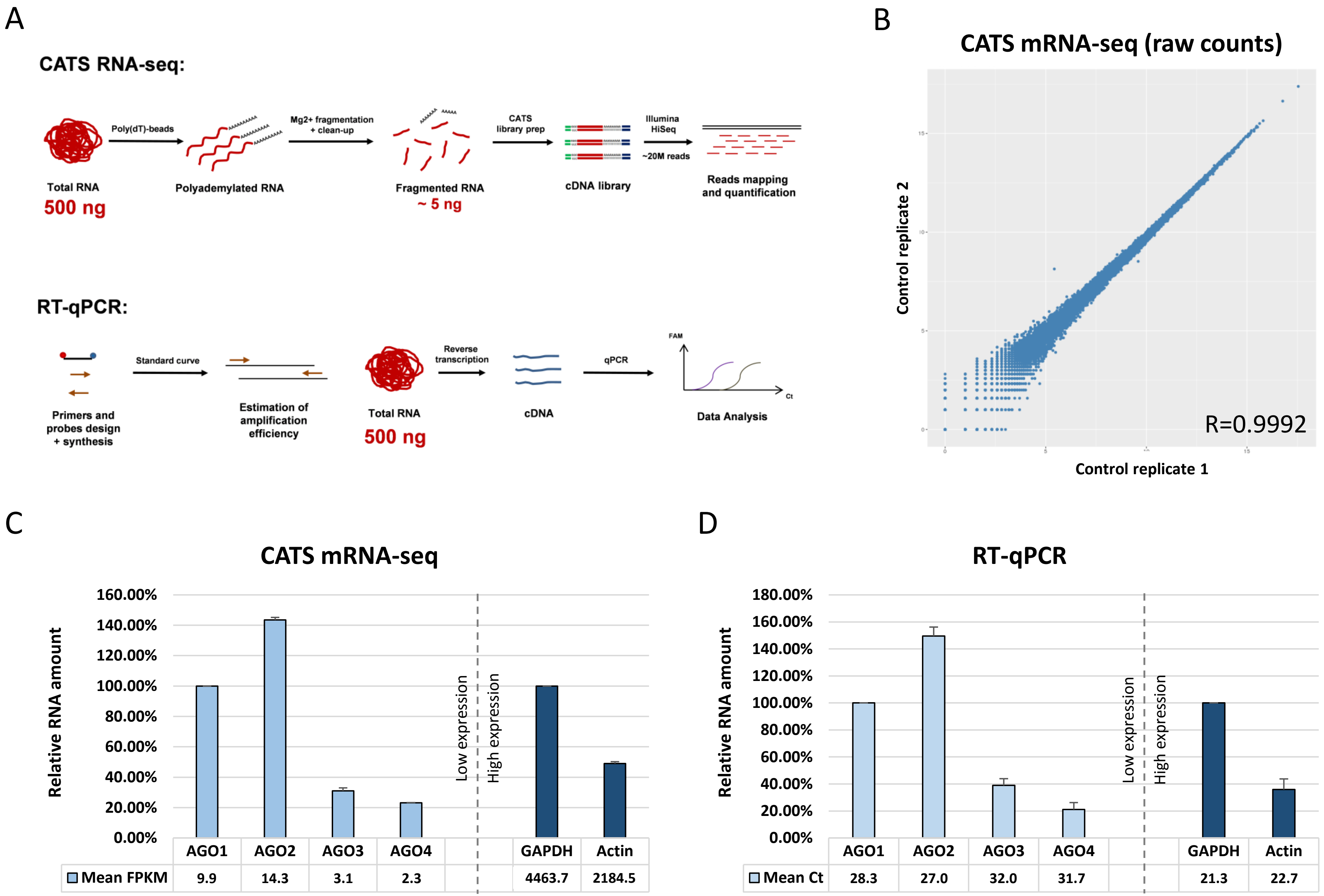

Supplementary Figure 3

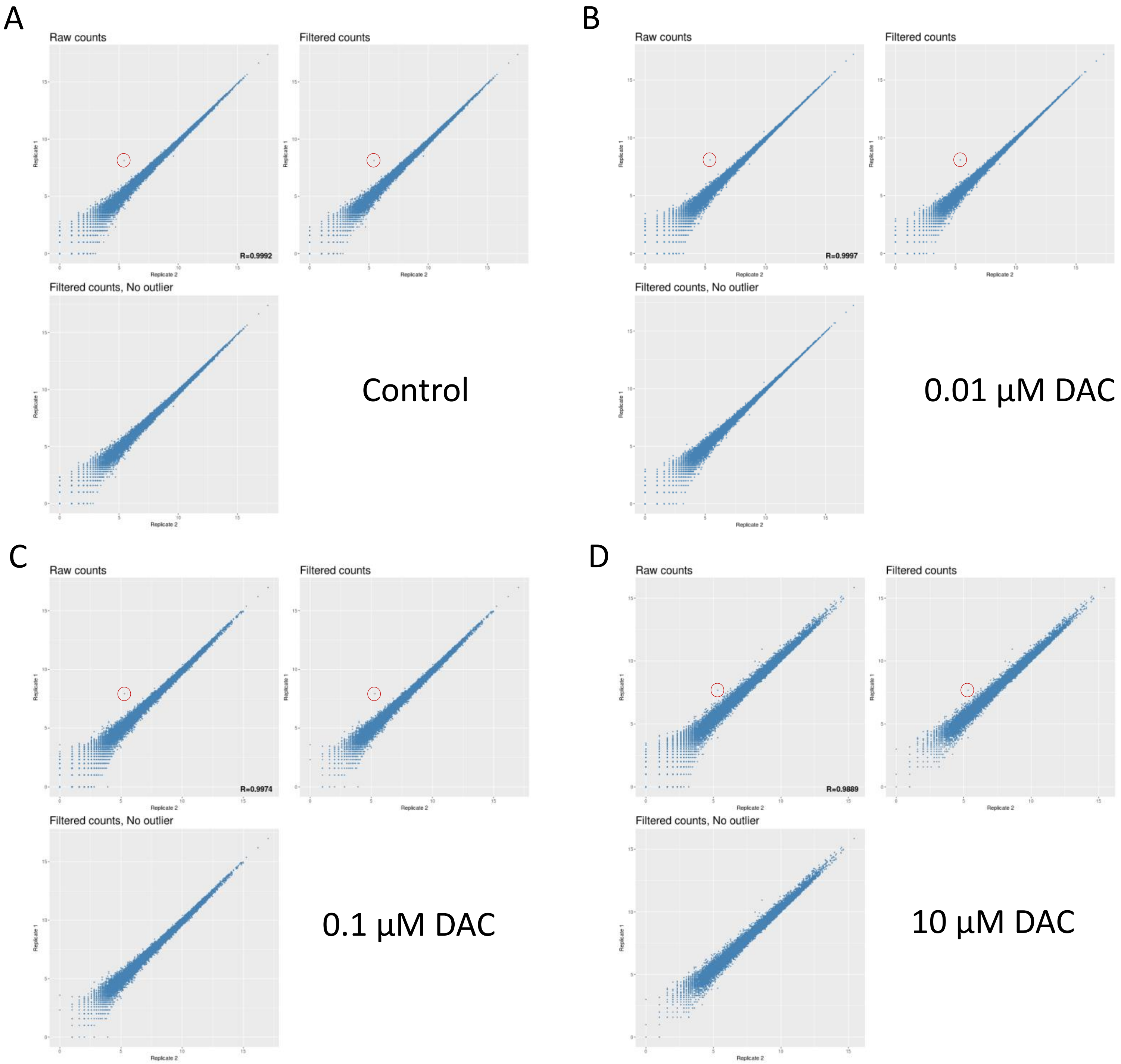

Supplementary Figure 4

A

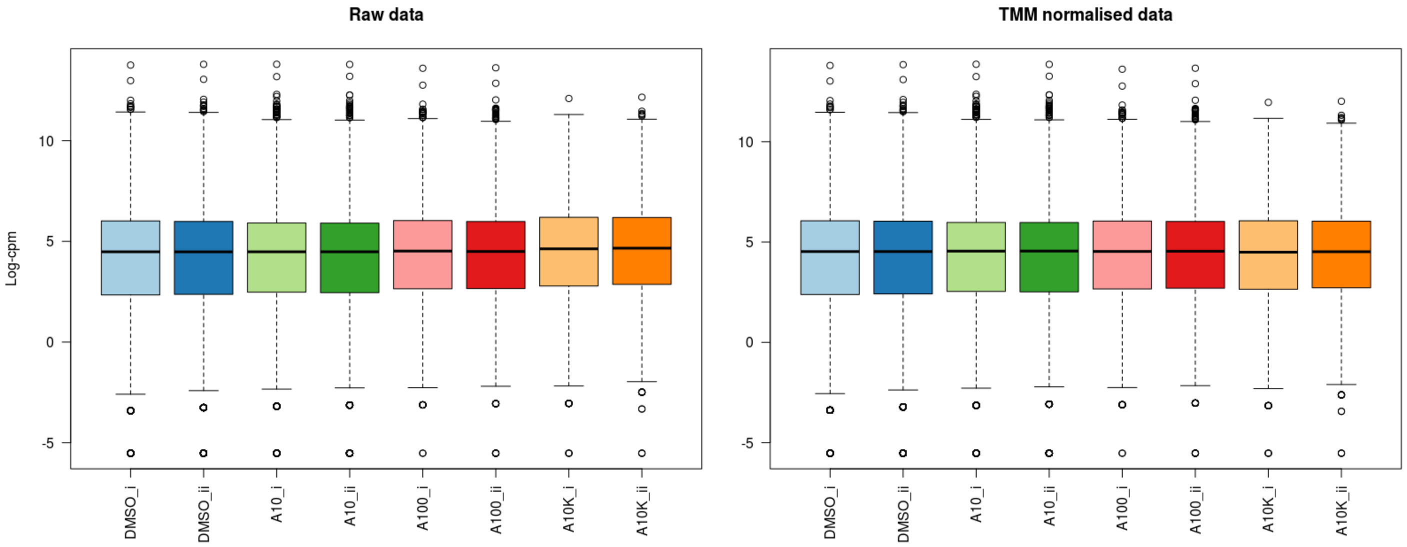


B

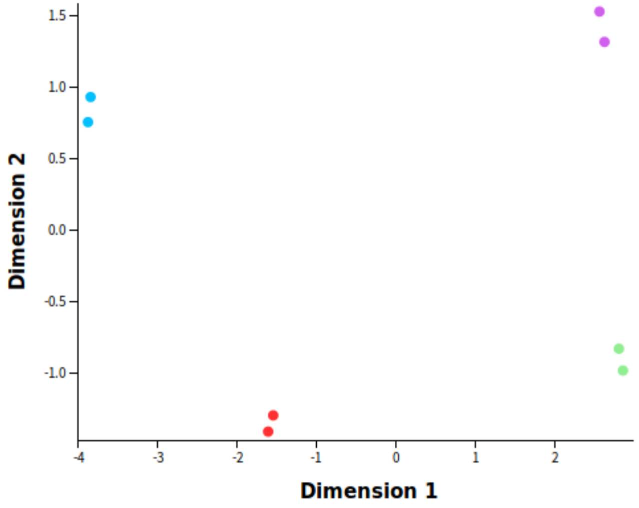

### Supplementary Figure 5

## A

#### Calculation of log-fold-change (LFC) by “FPKM method”

$FPKM^{geneA}$ = (RAW COUNTS $^{geneA}$ + 1) / TRANSCRIPT LENGTH $^{geneA}$ (in kilobases) * TOTAL MAPPED READS (in millions)

LFC = Log2 ($FPKM^{geneA}$)[control] - Log2 ($FPKM^{geneA}$)[DAC treated]

## B

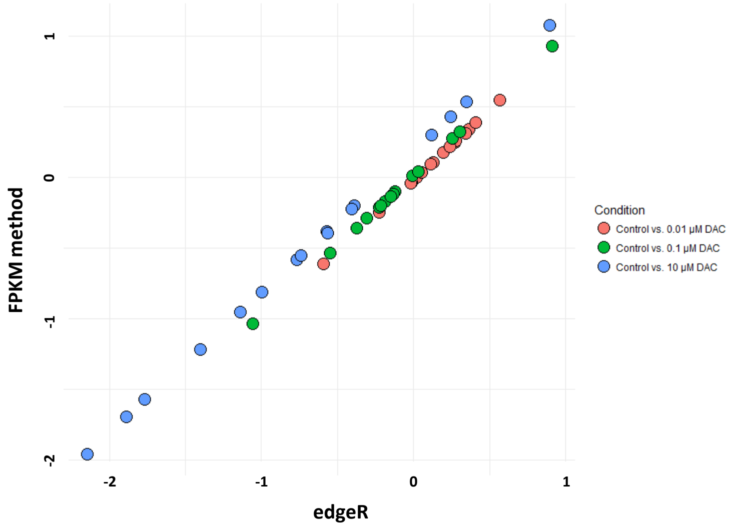


R = 0.9999

R = 0.9999

R = 0.9999

### Supplementary Figure 6

#### CATS mRNA-seq data processing

**Adapters trimming:**

```
cutadapt -u 3 raw.fastq | cutadapt -a AAAAAAAA - | cutadapt -a AAAAAAAN$ -a AAAAAAN$ -a AAAAAN$ - | cutadapt -a AGAGCACACGTCTG - | cutadapt -O 8 -g GTTCAGAGTTCTACAGTCCGACGATCNNN -o trimmmed.fastq –
cutadapt -m 20 trimmmed.fastq > SS.fastq
```

**Removing non-mRNA sequences:**

```
bowtie2-build trash.fa trash.fa
bowtie2 -q -p 12 -x trash.fa -U SS.fastq -S trash.sam
# trash.fa included the following human sequences: chrM, rRNA, tRNA, RN7S, RNU, snoRNA/scaRNA, VT-RNA, Y-RNA,
samtools view -b -@ 12 trash.sam -o trash.bam
bam2fastx --fastq -Q -o no_trash.fastq -N trash.bam
```

**Obtaining mRNA reads counts:**

```
bowtie2-build protein_coding_transcripts.fa protein_coding_transcripts.fa
bowtie2 -q -p 12 -x protein_coding_transcripts.fa -U no_trash.fastq -S no_trash.sam
samtools view -b -@ no_trash.sam -o no_trash.bam
express --f-stranded --no-bias-correct protein_coding_transcripts.fa no_trash.bam
```

#### CATS bisulfite-seq data processing

**Adapters trimming:**

```
cutadapt -a GATCGGAAGAGCACACG raw.fastq | cutadapt -e 0.2 -a A{100} - | cutadapt -a AGAGCACACGTCTG - | cutadapt -O 8 -g GTTCAGAGTTCTACAGTCCGACGATCGGG -o UKbis_trimmmed_new.fastq –
cutadapt -m 40 trimmed.fastq > SS.fastq
```

**Prepararion of Bismark reference genome:**

```
bismark_genome_preparation --bowtie2 --verbose link-to-folder-containing-human-reference-genome/Bismark
bismark --bowtie2 -n 1 -p 12 link-to-folder-containing-human-reference-genome/Bismark SS.fastq
```
